# Salient 40 Hz sounds probe affective aversion and neural excitability

**DOI:** 10.1101/2022.02.26.482077

**Authors:** F Schneefeld, K Doelling, S Marchesotti, S Schwartz, K Igloi, A-L Giraud, LH Arnal

## Abstract

The human auditory system is not equally reactive to all frequencies of the audible spectrum. Emotional and behavioral reactions to loud or aversive acoustic features can vary from one individual to another, to the point that some exhibit exaggerated or even pathological responses to certain sounds. The neural mechanisms underlying these interindividual differences remain unclear. Whether distinct aversion profiles map onto neural excitability at the individual level needs to be tested. Here, we measured behavioral and EEG responses to click trains (from 10 to 250 Hz, spanning the roughness and pitch perceptual ranges) to test the hypothesis that interindividual variability in aversion to rough sounds is reflected in neural response differences between participants. Linking subjective aversion to 40 Hz steady-state EEG responses, we demonstrate that participants experiencing enhanced aversion to roughness also show stronger neural responses to this attribute. Interestingly, this pattern also correlates with inter-individual anxiety levels, suggesting that this personality trait might interact with subjective sensitivity and neural excitability to these sounds. These results support the idea that 40 Hz sounds can probe the excitability of non-canonical auditory systems involved in exogenous salience processing and aversive responses at the individual level. By linking subjective aversion to neural excitability, 40 Hz sounds provide neuromarkers relevant to a variety of pathological conditions, such as those featuring enhanced emotional sensitivity (hyperacusis, anxiety) or aberrant neural responses at 40 Hz (autism, schizophrenia).

## Introduction

Beyond cultural variations in aesthetic tastes, there exists an important variability in the way individuals are sensitive and reactive to sounds, whether pleasant (e.g., baby laughs, ASMR) or unpleasant (e.g., screams, chalk on a blackboard). In extreme cases, sound hypersensitization can trigger pain (1), audiogenic seizures (2) or excessive stress sometimes resulting in aggressive reactions such as in the shaken baby syndrome (3). Given sound potential pathogenicity, it is critical to understand the neural mechanisms underlying such variability between individuals. Yet, the sonic features that drive hypersensitization and how they affect neural circuitry remain unclear.

Recent works have shown that alarm auditory signals (*e*.*g*., screams, sirens) exploit a restricted 30–80 Hz range of temporal modulations known as *roughness* to communicate danger, trigger negative affective responses and elicit rapid reactions (4). Unlike conventional noise, roughness can trigger aversive responses at relatively mild sound intensity (5). Therefore, it is particularly well-suited to investigate the neural underpinnings of auditory aversion in humans. Mapping behavioral and brain responses in the roughness range and beyond, intracranial EEG studies recently revealed that the non-linear pattern of aversion across frequencies strikingly resembles the spectral profile of a well-known auditory neural signal, the auditory steady-state response (ASSR) (5). This suggests that aversion to rough sounds might relate to the capacity of repetitive transients to induce sustained responses in the brain. However, subjective aversion profiles, though highly reliable across experiments when averaging data across participants, show substantial variability across participants. If the link between aversion and brain responses holds at the individual level, then we should expect similar variability in the ASSR signals of our participants. Here, we primarily sought to explore, using non-invasive surface EEG, whether aversive sound rating patterns relate to neural excitability at the individual level. More specifically, we tested the hypothesis that increased aversion to roughness could be underpinned by enhanced auditory steady-state responsiveness to sounds.

Electrophysiological brain signals are known to maximally entrain in a sustained manner to repetitive acoustic transients at 40 Hz. Since the study by Galambos and colleagues (6), a wealth of experimental work has investigated this phenomenon. Although often used as an index of early auditory process integrity (7), it has been hypothesized that these signals more generally index long-range thalamo-cortical synchronization and information transfer in the brain (8, 9). The 40-Hz ASSR is often considered a promising biomarker of various neuropsychiatric (10, 11), neurodevelopmental (12, 13) and neurodegenerative disorders (14). However, the link between these disorders and auditory processing is unclear. While some disorders are comorbid with hearing problems (15, 16), the prevalence of ASSR impairments in these diseases might point to dysfunctions beyond or outside classical auditory pathways. Indeed, whereas previous EEG studies a priori locate the source of ASSR generators in the auditory cortex, recent findings using alternative reconstruction algorithms or spatially resolved intracranial recordings suggest that rough sounds additionally synchronize non-canonical auditory systems involved in salience and aversion processing (5, 17, 18). Therefore, impaired ASSR responses –as observed in neuropsychiatric disorders– might actually reflect dysfunctional sensory and emotional processing of exogenous stimulation in these networks.

To summarize, previous findings demonstrate (*i*) that 40 Hz sounds are perceived as aversive, (*ii*) that they recruit canonical and non-canonical auditory networks involved in aversion processing, and (*iii*) that the neural processing of these sounds might be altered in a variety of brain disorders featuring abnormal emotional responses. These converging observations suggest that the neural excitability of aversion networks –as revealed by the strength of ASSR– might influence emotional reactivity to these sounds, as well as indicate abnormal excitability in neuropsychiatric disorders. Here, we aimed to clarify the link between emotional reactivity and neural excitability to rough sound at the individual level in the normal population. In line with previous observations that ASSR are compromised in neuropsychiatric and neurodegenerative diseases classically featuring enhanced anxiety, we further tested how these responses interact with individual anxiety level.

## Methods

### Stimuli

All stimuli were digitally generated using MATLAB with a sampling rate of 96 kHz and presented in a pseudo-randomised order using Psychtoolbox (Version 3.0.12). Click trains (1 s duration, with 100 ms sine ramping onset and offset; click rise/fall time of 0 ms, plateau time of 1 ms, presented at ~60 dB SPL at frequencies varying between 10 and 250 Hz).

### Paradigm

The first behavioral experiment (Exp. 1, pilot, N=12, 6 females) measured subjective aversion as a function of the frequency of click trains (10, 20, 40, 60, 80, 100, 120, 140, 170, 200 and 250 Hz). After the presentation of each stimulus, participants reported their aversion to click trains of varying frequencies on a 5-level Likert-like subjective scale ranging from tolerable (1) to unbearable (5). The second experiment (Exp. 2, N=38, 25 females) replicated Exp. 1 while recording participants’ EEG signals. In Exp. 2, stimuli (click trains at 10, 20, 30, 40, 60, 90, 130, 180 and 250 Hz) were presented in pairs, separated by a period varying from 800 to 1200 ms. Participants were required to rate the unpleasantness of the second stimulus of the pair only, to avoid memory effects, on an analog slider scale ranging from 1 (tolerable) to 10 (unbearable). Pairs were made of either the same (repeated, 80% of all pairs) or different (oddballs, 20% of all pairs) stimuli. Only pairs containing repeated stimuli were analyzed in the current study. All (oddball and repeated) pairs were presented in pseudorandomized order. Two participants did not complete the task correctly and were removed from subsequent analyses.

### Behavioral data analysis

Individual aversion profiles in Exp. 1 and Exp. 2 were obtained by averaging aversion scores per stimulus frequency, which were subsequently z-scored per participant to account for baseline individual aversion biases (inter-individual variability of overall sound aversion across all frequencies). Qualitative inspection of individual profiles suggests the existence of two distinct aversion profiles: on the one hand, *roughness* responders exhibiting a non-linear response with heightened aversion to low frequencies (*low, 30–60 Hz*), and on the other hand, *pitch* responders, in whom aversion follows a linear profile with a maximal response for the highest frequencies (*high*, >130 Hz; Fig. 1). To quantify this dual patterning of aversion profiles across participants –namely, to test if those participants sensitive to high frequencies were less sensitive to low frequencies and conversely–, we computed the correlation of aversion between each tested frequency across participants. Computing the sign of the difference between aversion scores in *low* [30–60 Hz] minus *high* [130–250 Hz] frequency ranges, we then categorized participants as *roughness* responders (*low* – *high*>0) or *pitch* responders (*low* – *high*<0).

**Figure 1:**
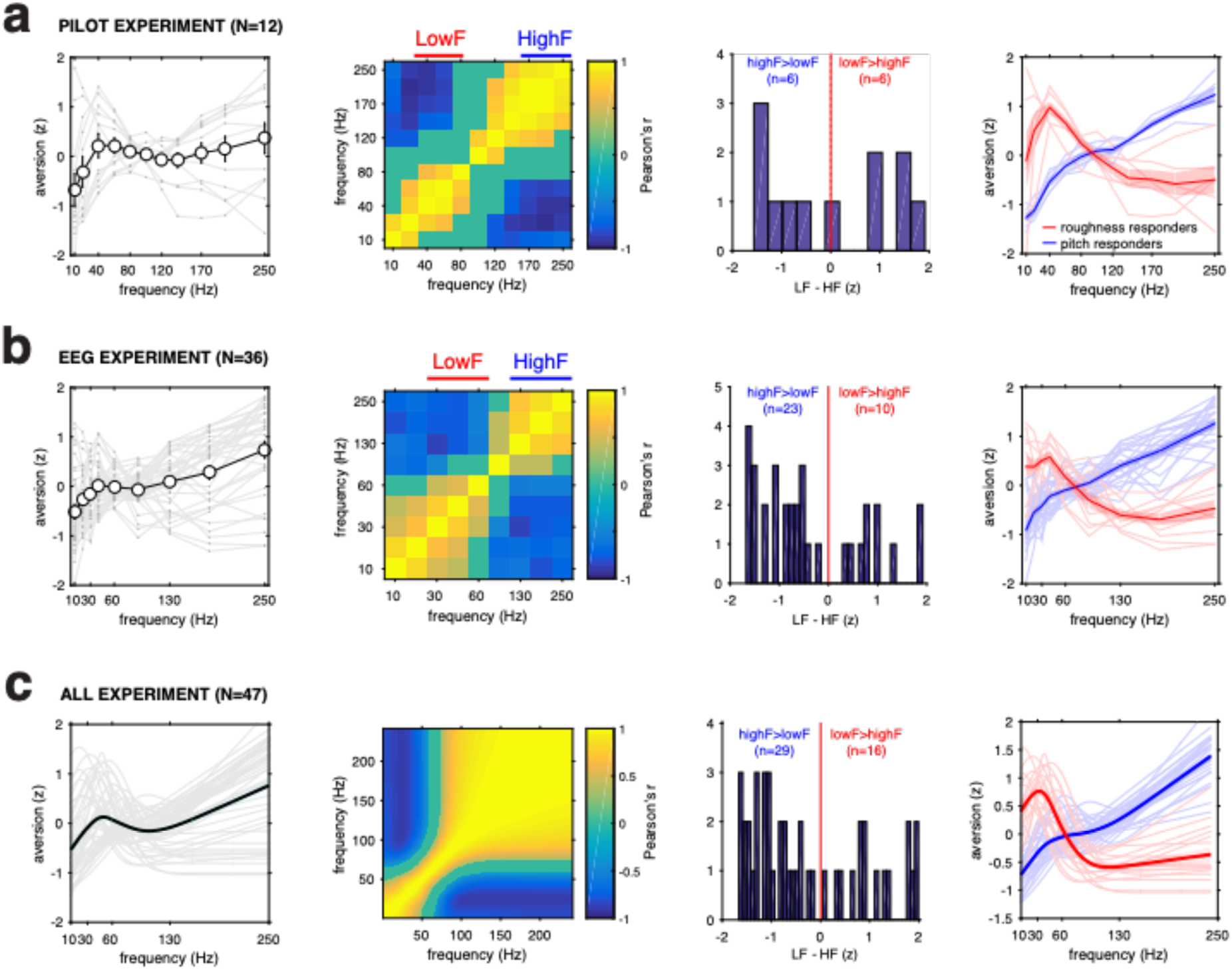
Behavioral aversion ratings. **a**. Left panel: Pilot experiment: behavioral aversion ratings as a function of stimulus frequency. White circles and error bars represent mean across participants +/- SEM. Grey lines correspond to individual aversion curves. Middle left: correlation of aversion scores across frequencies and individuals. Positive correlation values (yellow) show that interindividual variability is preserved across neighboring frequencies. Negative correlation values (blue) show that participants who rate low frequencies (30–60 Hz, *low*) as more aversive rate high frequencies (130– 250 Hz, *high*) as less aversive, and reciprocally, evidencing a distinction between *low* and *high* responders. Middle right: determination of *low* minus *high* responder groups: participants with positive *low* minus *high* aversion values are categorized as *low* responders and reciprocally for *high* responders. Right panel: individual aversion rating curves categorized as *low* (red) and *high* (blue) responders. **b**. Same as in (a.) for the EEG experiment. **c**. Same as (a.) and (b.) grouping interpolated data from the pilot and the EEG experiment (see supplementary information).

To check the replicability of aversion profiles between Exp. 1 and Exp. 2 and to be able to subsequently pool the results across experiments, we then applied the following modelling procedure. As we selected slightly different stimulus frequencies (in a similar frequency range) for Exp.1 and Exp. 2, we interpolated the data at each frequency in the selected frequency range. Stemming from the hypothesis that non-linear profiles result from the differential weighting of non-linear *roughness* and *pitch* profiles (Figure 1), we then sought to fit individual behavioral data aversion scores. In each participant, we fit a summation between a gaussian distribution and a linear model (Supplementary Figure 1 and the equation below):

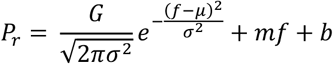

Where *P*_*r*_ is the predicted rating of a given trial and *f* is the stimulus frequency of that trial. The model contains 5 parameters: 3 for the gaussian shape, a mean *µ*, a standard deviation *σ* and a gain *G* to normalize the height of the peak to fit in the scale of the subjective ratings; and 2 parameters for the linear portion, slope *m* and intercept *b*. The parameters were fit by minimizing the squared error between the model predictions and the true sound rating taking the stimulation frequency as input. Minimization was performed using the fminsearch function in MATLAB. The model allowed us to interpolate our results to untested frequencies and better compare across experiments using different stimulus frequencies.

### EEG analyses

EEG recording was performed using the DSI 24 EEG headset (Wearable Sensing, San Diego, USA). This headset is equipped with 21 dry electrodes, placed according to the 10/20 international standard and permits wireless data transmission at a 300 Hz sampling rate. Two additional reference electrodes were placed at the ears. The data was common-average re-referenced and chunked in 1500 ms epochs (500 ms before to 1000 ms after the click train onset). For each trial, we then converted the data into z-scores using the pre-stimulus baseline activity (from 200 ms before to stimulus onset) for each electrode.

All EEG data analyses were performed in MATLAB (MathWorks) using the Fieldtrip (http://fieldtrip.fcdonders.nl) package and in-house custom code. Trials were visually inspected, and those with obvious artifacts were removed. An independent component analysis as implemented in FieldTrip was used to correct for eye blinks and eye movements. Adding to the two participants who did not perform the task correctly, three participants were excluded from the EEG analysis due to poor EEG signal quality, leaving thirty-three participants to be included in EEG analyses. Auditory ERPs and ASSRs were calculated for each individual, electrode and stimulus frequency. For ASSRs, we measured the power of neural responses filtered at the stimulus frequency. Amplitude time series were computed by band-pass filtering the EEG signal at the stimulus frequency F ± 0.5 Hz. The envelope of each narrow-band signal was obtained by taking the absolute value of the analytic signal resulting from a Hilbert transform. ASSR values were baseline-corrected by subtracting the power of the EEG signal at stimulus frequency during a 200-ms period preceding sound onset to capture endogenous activity. We then transformed these time-courses into z-scores per trial and electrode and participant. Given that the absolute magnitude of auditory responses (ERPs) substantially varied across participants (whether for SNR or physiological grounds), we quantified ASSR relative to the canonical auditory ERPs to *high* [130–250 Hz] sounds. We selected these frequencies because ASSR power is negligeable and minimally interacts with ERP components at these frequencies. As our goal was to explore and relate inter-individual brain responses to a variety of measurements (subjective aversion, anxiety) and given the relative SNR disparity across participants, we standardized ASSR responses in each participant as follows. We subtracted early evoked response to high frequencies (value of the N1 in the 3 central sensors) to the ASSRs at each frequency and electrode. Reasoning that *roughness* responders should exhibit larger responses to *low* than *high* sounds (and conversely for *pitch* responders), this procedure allowed standardizing these measurements across participants, and provides a relative metric to assess *low* versus *high* excitability at the individual level. Finally, to avoid a potential contamination of entrained ASSR by ERPs to sound onsets (*e*.*g*., N1) and by motor preparation responses, we focused our analysis on a time window—defined by data visual inspection of the data— from 200 to 600 ms after stimulus onset.

### Correction for Multiple Comparisons

To assess the statistical difference between the experimental conditions while controlling for multiple comparisons, we performed nonparametric cluster analyses (19) for each statistical test. This method is based on comparing the cluster statistic of adjacent timepoints (significant at p<0.05) to the same statistic ran on randomly permuted data. The nonparametric statistic was performed by repeating 1000 times the calculation of a permutation test where the experimental conditions are randomly intermixed within each subject. For each of these permutations, we extracted the maximum (absolute value) cluster-level statistic. Finally, we calculated the corrected p-values by comparing the cluster-level statistics of our original data with those of all permutations. All the results presented are corrected for multiple comparisons using this method unless otherwise stated.

## Results

### Subjective aversion: roughness versus pitch responders

To investigate the inter-individual variability or subjective aversion across frequencies, we first analyzed the data from a previous behavioral experiment (Experiment 1, see (5). Twelve participants listened to a series of 1-s click trains of varying click frequency (ranging from 10 to 250 Hz) and rated the sounds’ aversiveness from 1 (tolerable) to 5 (unbearable). Figure 1a (left panel) shows that aversion varied in a non-linear, non-monotonic manner with increasing sound rates. Qualitatively, individual rating profiles appeared quite variable across participants, some exhibiting higher aversion to *low* than *high* frequencies and others exhibiting the inverse pattern. To quantify this preliminary observation, we tested the correlation of ratings across participants and frequencies (Figure 1a, middle left panel). We observed that while aversion patterns were very similar across participants between adjacent frequencies (along the diagonal of the correlation matrix), aversion patterns were negatively correlated between low (<80 Hz) and high frequencies (>130 Hz). This effect reveals a dual pattern of auditory aversion across participants: while a subset of participants was most sensitive *to high* frequencies and relatively less to *low* (*pitch* responders, N=6), the other half (*roughness* responders, N=6) was more sensitive to lower, temporally salient frequencies but not to high frequencies. To categorize participants as *roughness* or *pitch* responders (Figure 1a, middle right panel), we measured the difference between averaged ratings of *low* (30–60 Hz) and *high* (130–250 Hz) sounds. This allowed us to assign each participant to either a *roughness* responders (*low* minus *high* > 0) or a *pitch* responders (*low* minus *high* < 0) group. Figure 1a (rightmost panel) reproduces Figure 1a (leftmost panel) color coded as a function of *roughness* (red) versus *pitch* (blue) responder groups.

To replicate these behavioral results and investigate their underlying neural correlates, we repeated the same experiment (Experiment 2) using slightly different stimulus rates (ranging from 10 to 250 Hz) in 33 participants while recording electroencephalographic (EEG) responses to these sounds. Focusing on behavioral data, we found that the pattern of ratings and ensuing analyses (Figure 1b) replicated the results obtained in Exp. 1, although the ratio of *pitch* (N=23) versus *roughness* (N=10) responders was slightly less balanced (Figure 1b). Finally, to pool the data from both experiments for subsequent analyses, we developed a non-linear curve fitting procedure (see Methods and supplementary data section) which reproduced the dual pattern of *roughness* versus *pitch* responsiveness to aversion (Figure 1c). To test the replicability of this non-linear pattern between experiments 1 and 2, we measured the Pearson’s correlation between interpolated aversion profiles (averaged across participants within each experiment) and confirmed that these profiles were highly similar between the two experiments (Pearson’s r = 0.79, p = 1e-53).

### Event related potentials relate to subjective aversion

To analyze brain responses to click trains of varying rates, we first measured the magnitude of ERPs across channels and participants (Figure 2a). This analysis replicated most features of the main results previously observed in intracranial recordings (5). Focusing on the early, N1 component, we observed that the magnitude of this component linearly increased as a function of click trains frequency (Pearson’s r=-0.91, p= 0.0007). Measuring the amplitude of ERPs in a late [200–600 ms] time-window, we observed that brain responses to low frequencies (low, 30–60 Hz) were more negative than responses to higher (*high*, >130 Hz) frequencies (t(32)=9.22, p=0.01) with a profile that qualitatively resembled the non-linear aversion rating profiles.

**Figure 2:**
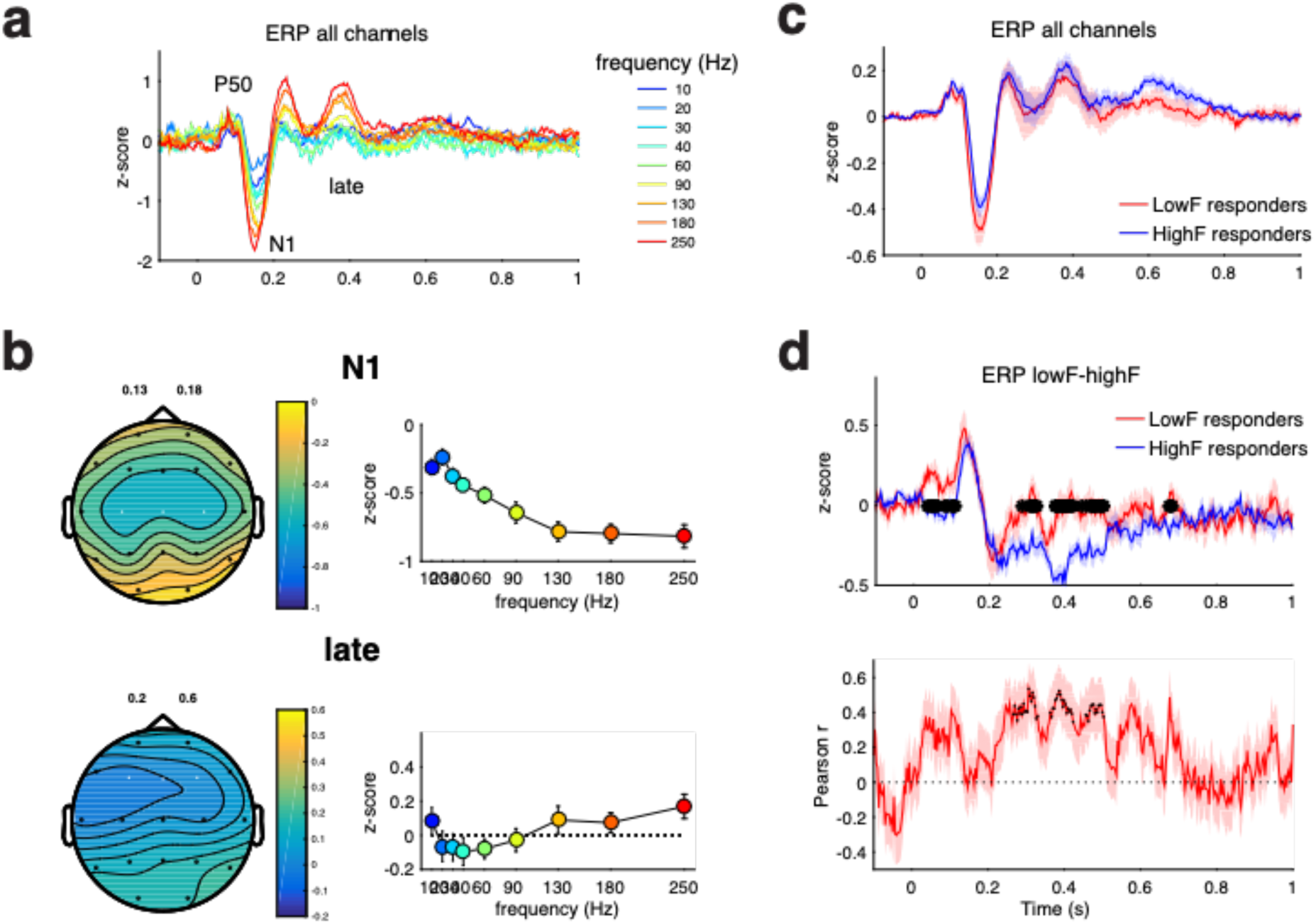
ERP responses across frequencies. **a**. ERP responses across time and averaged across all EEG channels as a function of stimulus frequency. **b**. Top left panel: Topography of the N1 response. Top right panel: N1 magnitude as a function of stimulus frequency. Bottom left panel: Topography of the late [0.2 – 0.6 s] responses. Bottom right panel: late response magnitude as a function of stimulus frequency. Selected channels (exhibiting maximal magnitude) are highlighted in white **c**. ERP responses across frequencies do not differ between *Low* (red) and *high* (blue) responders. **d**. Top panel: difference between low [30-60Hz] and high [130-250 Hz] frequency ERPs for *low* (red) and *high* (blue) responders. Stars indicate significant (p<.05 corrected) statistical difference. Bottom panel: Correlation between aversion ratings and *low* minus *high* ERPs. black dots indicate (cluster corrected) statistical significance at p<.05 level.

We first compared the magnitude of ERPs across all stimulus frequencies between the two groups of participants over central N1 channels (Figure 2c). This analysis did not yield significant differences between groups, suggesting similar levels of activity in responses to click trains. However, our primary hypothesis was that *roughness* and *pitch* responders might exhibit different neural responsiveness to *low* versus *high* frequencies, respectively. We therefore sought to compare the relative ERP magnitude to *low* versus *high* stimulus modulation frequencies between the two groups. To do so, we first computed the difference between *low* and *high* ERPs in each group (Figure 2d). This analysis revealed significant differences in early components (P50) as well as several significant clusters in the late [0.2– 0.6 s] time-window. We then tested the Pearson’s correlation between individual aversion ratings and the *low* minus *high* ERP difference across time, and found that the correlation was maximal and significant in the late time-window (Pearson’s correlation r_[200–600 ms]_ =0.48, p= 0.004).

### Auditory Steady-State Responses to low sounds reflect subjective aversion

*Based on* the hypothesis that temporally salient *low* sounds should entrain brain responses in a steady-state, sustained manner, we measured the power of steady-state responses (ASSR) at the rate of the exogenous stimulation in the late time window (restricted to [200–600 ms] to avoid potential contamination by onset and offset responses). ASSR power averaged across all electrodes showed a sustained response throughout the sound duration (Figure 3a, left panel). Focusing on brain responses to low sounds (30–60 Hz) across all participants (both groups together), we observed that ASSR power in this range was maximal in central and anterior contacts, a topographical pattern that is compatible with the recruitment of auditory and frontal regions as observed in intracranial recordings in humans (5, 18) and murine models (20).

**Figure 3:**
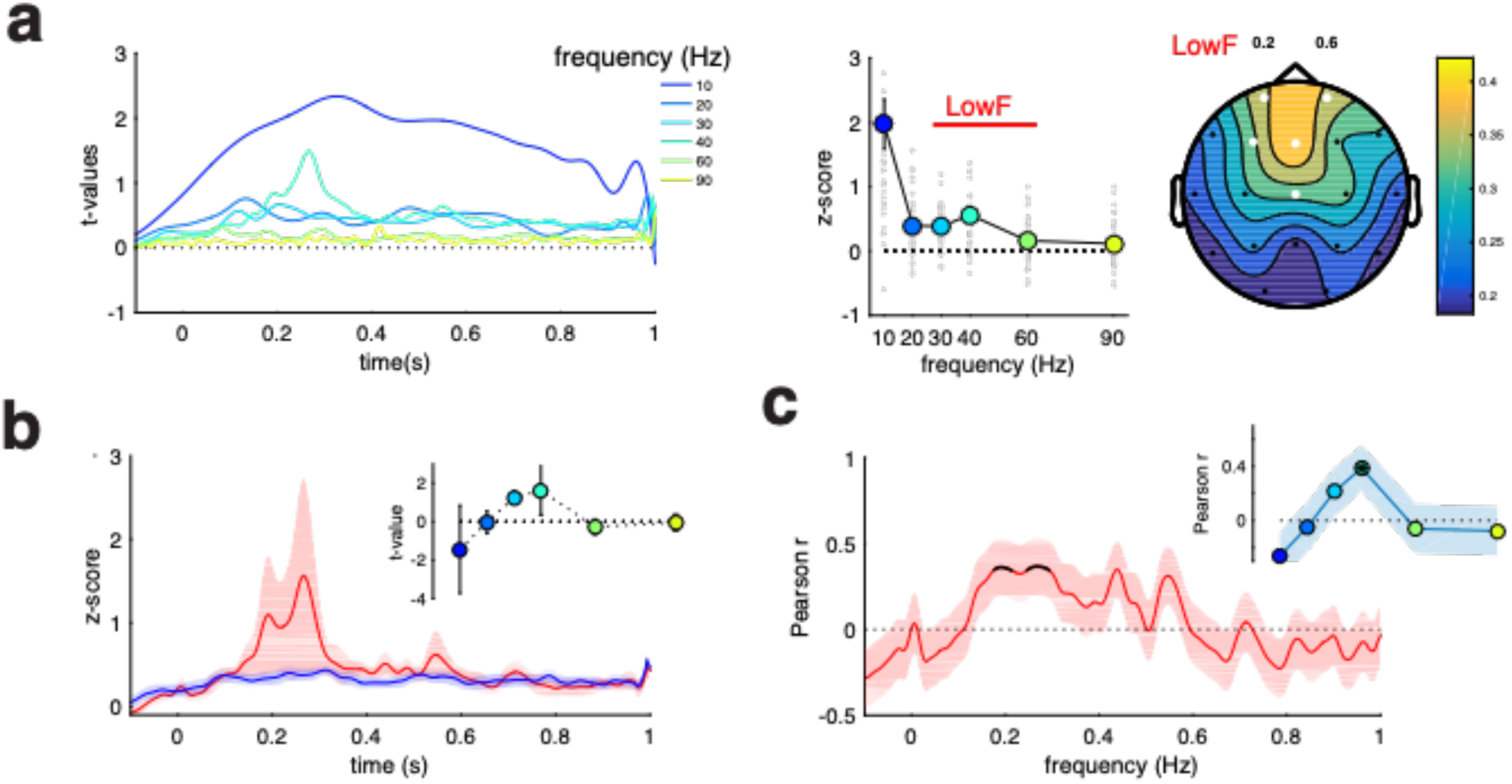
Time-frequency analysis of low steady-state responses (ASSR). **a**. Left: time course of ASSR responses at each stimulation frequency. Middle: topography of *Low* (30–60 Hz) ASSR power averaged over the late [.2–.6 s] time-window. Right: ASSR power per stimulation frequency averaged across the late time-window. Grey dots represent individual data points. **b**. Time-course of *low* ASSR power for low (red) and high (blue) responders. Upper right inset: averaged ASSR power difference between *low* and *high* responders in the late-time time window. **c**. Time-course of Pearson’s correlation between *low* ASSR power and aversion ratings. Black lines represent significant correlation values at p<.05 level (cluster corrected). Upper right inset: Pearson’s correlation r-values per stimulation frequency in the late-time time window. Star indicates significance at p<.05.

We then aimed to assess the difference in ASSRs for *roughness* and *pitch* responders in central-anterior sensors (Figure 3b). Despite the qualitative difference between *roughness* and *pitch* responders, this contrast did not reach statistical significance. We then tested the correlation in time between *low* ASSRs and individual subjective ratings in the same set of electrodes and identified several correlation clusters in the late window. Averaged over the late time-window, the correlation proved to be maximal and significant at 40 Hz (Figure 3c).

### Anxiety increases aversion and ASSRs to low frequency sounds

We then tested the hypothesis that anxiety might interact with the excitability of aversion networks. We tested the relationship between individual anxiety levels (as assessed using the STAI questionnaire) and (*i*) subjective aversion and (*ii*) brain responsiveness to *low* sounds. Focusing on subjective ratings from Exp. 1, we first determined that aversion ratings significantly correlated with Trait-anxiety scores (Figure 4a). In other words, while the most anxious participants rated rough, *low* (30-60 Hz) sounds as more unpleasant, the least anxious participants rated louder, *high* trains (>180 Hz) more negatively. Replicating this analysis in Exp. 2, we found a similar pattern of correlation with STAI-T scores (Pearson correlation between Exp. 1 and 2 correlation profiles: r = 0.929; p = 0.001), although non-significant. In addition, pooling interpolated data from Exp. 1 and 2 (see Fig. 1c) confirmed that across all participants and experiments, trait anxiety positively correlated with subjective aversion to *low* and negatively correlated with *high* sounds.

**Figure 4:**
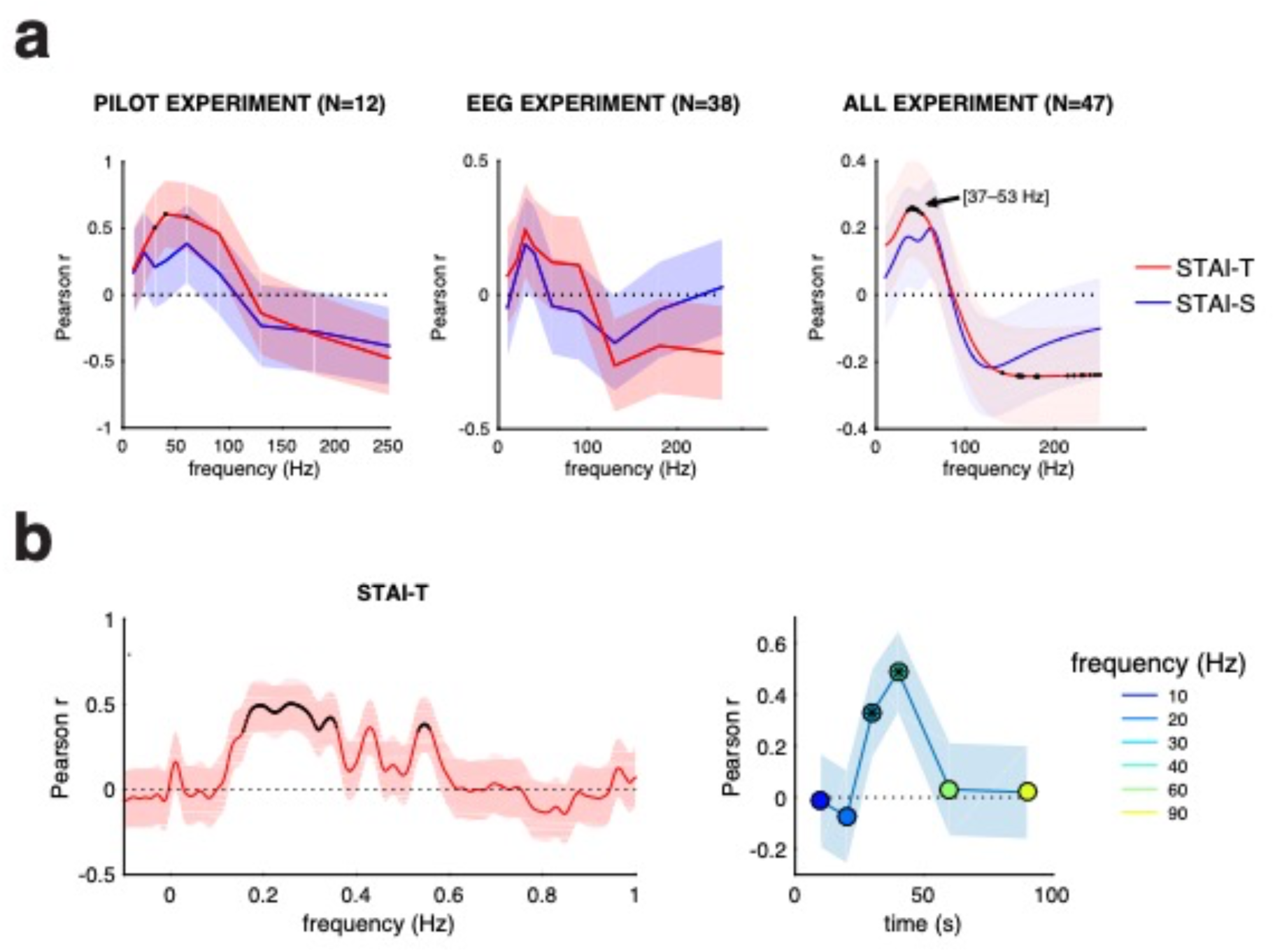
Relationship between auditory aversion and anxiety at the inter-individual level **a**. Pearson’s correlation between aversion ratings and individual anxiety levels assessed via STAI questionnaires for STAI-Trait (red) and STAI-State (blue). Shaded areas represent SEM of the correlation. Left: pilot experiment. Middle: EEG experiment. Right: same as in left and middle panels grouping interpolated data from the pilot and the EEG experiment (see supplementary information). Time-course of Pearson’s correlation between *Low* ASSR power and anxiety STAIT-T scores. Black lines represent significant correlation values at p<.05 level (cluster corrected). Right: Pearson’s correlation r-values per stimulation frequency in the late-time time window. Star indicates significance at p<.05. Right: topography of correlation r-values between *Low* ASSR power and anxiety STAIT-T scores.

Focusing on ASSR responses to *low* sounds, we then measured the relationship between interindividual anxiety (STAI-T) scores and cerebral responsiveness to aversive sounds. We observed that whereas anxiety does not interact with early auditory ERPs, this analysis revealed a significant correlation between anxiety and strength of late [200-600 ms] sustained auditory steady-state responses (ASSR) to low sounds (Fig. 4b). Focusing our analysis of ASSR in the late time-window, we found that the correlation was maximal and significant only at 30 and 40 Hz.

## Discussion

### Aversion to roughness links to steady-state excitability

Here, we investigated the relationship between individual subjective aversion and neural responsiveness. Replicating our previous work (5), we show that aversion to repetitive sounds in the roughness range and beyond is not a linear function of stimulus frequency. When averaged across participants, aversion follows a non-linear pattern, peaking at 40 Hz in the roughness range [30–80 Hz] and linearly increasing as a function of stimulus frequency above 130 Hz. However, further investigating inter-individual variability, we found that although highly reliable across experimental replications (see (5) and current study), this pattern substantially varied across participants. While some participants’ aversion profile followed a linear pattern seemingly determined by stimulus frequency (here proportional to sound intensity), other participants’ aversion profile revealed a higher sensitivity to sounds in the roughness range, most prominent at 40 Hz. Exploring the variability of individual aversion profiles, we discovered that some participants were maximally sensitive to high frequency sounds (*pitch* responders), others to low frequencies (*roughness* responders) or exhibited a composite pattern between these two extremes.

When studying the neurophysiological origin of the non-linear pattern of aversion across frequencies using intracranial recordings in epileptic patients, we recently suggested that unlike aversion to higher frequencies in the pitch range (>130 Hz), aversion to rough frequencies is best accounted for by the sustained entrainment of activity of brain regions involved in salience and aversion processing (5). In other words, beyond triggering classical onset related potentials (ERPs) in the auditory cortex, the non-linear effect of subjective aversion in the roughness range is arguably induced by the additional steady-state neural entrainment (ASSR) of widespread non-canonical auditory networks (5, 17, 18).

Considering the disparity of aversion profiles between participants, we then hypothesized that one possible origin of this inter-individual variability might be the relative excitability of these networks and related ASSR responses across participants. We predicted that individual sensitivity to roughness (versus stimulus frequency) might be determined by the relative magnitude of ASSR and ERP signals, respectively. Relating these EEG signals to individual aversion profiles across 33 participants, we found that aversion to rough (*low*) sounds correlated with the magnitude of ASSR signals (relative to *high* ERP magnitude). This supports our initial prediction, namely that sensitivity to rough sounds is linked to neural steady-state responsiveness, at the individual level. In light of the notion that ASSRs reflect the entrainment of widespread, non-canonical neural networks involved in salience processing, this finding suggests that the relative aversion to rough sounds, in particular at 40 Hz, might provide a valuable proxy of inter-individual excitability in those networks.

### Anatomical and functional sources of ASSRs and related aversion

The phenomenological distinction between roughness and pitch is commonly attributed to a dual coding strategy of temporal and spectral information in the canonical auditory pathway (21–23). However, recent results suggest that these two response modes may reflect the recruitment of distinct neural routes for sound processing. While pitch sounds are processed along the canonical auditory pathway, rough sounds may additionally target an alternative, non-canonical pathway ultimately targeting medial cortical and limbic regions ^5,16,17^. Here, we hypothesize that the inter-individual variability in neural and emotional responses to 40 Hz sounds might be accounted for by increased excitability of these non-canonical networks relative to canonical auditory pathways.

The anatomical origin and trajectory of such pathways, and how they trigger negative emotional responses remain elusive. A recent stream of findings in mouse models has shown that 40 Hz sounds entrain neural activity in medial temporal and limbic –not primarily auditory– regions notably involved in arousal, emotional and attentional processing ^19,24^. Interestingly, rough sounds transiently interrupt microsaccadic rhythms, suggesting that they impact oculomotor processes via non-primary auditory brainstem nuclei. That these sounds interact with non-auditory attentional processes concurs with other observations highlighting their impact on aversion (5), vigilance state and arousal (9, 24). This further highlights the propensity of rough sounds to affect early arousal and low-level emotional processes in a way that possibly prescinds higher-order auditory functions. Although more experimental work is needed to decipher the circuits involved and their influence on affective responses, the current results support the idea of an additional, non-canonical auditory circuit – although possibly overlapping with classical ones– involved in the affective, non-linear processing of roughness.

### Mapping between affective and neural responsiveness to rough sounds

Although brain responses to 40 Hz sounds are widely studied in neuroscience and frequently used in audiological assessment (25), why and how these sounds affect arousal and induce negative emotional responses remains neglected.

It has been known for decades that steady-state EEG responses to 40 Hz sounds are affected in various brain disorders, whether neuropsychiatric (10, 11), neurodevelopmental (12, 13) or neurodegenerative (14, 26, 27). Interestingly, many of these pathologies often feature (*i*) deficits in salience processing and arousal (28) and (*ii*) excitation/inhibition imbalance in medial temporal, limbic and salience-related networks (29–32). Many of these disorders further evidence atypical affective responses and are often associated with increased anxiety (33–36). However, the link between neural excitability and emotional reactivity to these sounds – whether in neuropsychiatric or neurotypical populations – remains unclear.

Anxious people often exhibit exaggerated responses to salient or unpleasant sensory events (1). Here, we show that interindividual sensitivity to *low* sounds (relative to *high* ones) correlates both with neural responsiveness to these sounds and with individual anxiety trait. In other words, the most anxious participants are more averse and show stronger neural entrainment to rough, *low* sounds than participants whose sound aversion is mostly driven by the frequency (here confounded with intensity) of the sounds. Considering recent works suggesting that 40 Hz sounds entrain non-canonical networks involved in aversion processing (5), these results suggest that these networks are more excitable in anxious participants, who in turn perceive these sounds as more unpleasant. As these effects more tightly link neural and emotional responsiveness to anxiety trait rather than state, one could conjecture that such systems are overall more excitable in anxious participants rather than being over activated at the time of the experiment.

Altogether, these results support the prediction that emotional and electrophysiological responses to 40 Hz sounds concurrently probe responsiveness of aversion networks and participants’ anxiety at the inter-individual level. The neuroanatomical and functional origin of such networks remains to be clarified e.g., in animal or pharmacological models focusing on emotional reactivity to these sounds. However, adding to previous works showing aberrant 40 Hz ASSRs in a variety of neuropsychiatric disorders, our results point to the possibility that anxiety implicates –and can be probed by– the over-responsiveness of brain networks involved in aversion to rough sounds. To conclude, the current work shows that aversion to 40 Hz sounds and ensuing brain responses constitute viable biomarkers to probe emotional responsiveness and neural excitability in non-canonical auditory networks. Such neuromarkers open new perspectives to assess the effect of therapeutic interventions on neural and emotional responsiveness at the individual level, in a variety of brain disorders featuring excessive emotional reactivity or neural responsiveness to temporally salient, rough stimulation.

## Acknowledgements

We thank Arnaud Desvachez and Blanca Marin Bosch for their contribution to experimental work. This work was funded by Swiss National Science Foundation project grant 163040 (A.G.), Fondation Pour l’Audition FPA RD-2020-10 (L.A.). *Author contributions:* L.H.A designed the experiments. F.S and L.H.A. collected the data. K.D. and L.H.A. performed the analysis. L.H.A. drafted the manuscript. F.S., K.D., S.M., S.S., K.I., A.L.G and L.H.A. corrected and approved the manuscript. *Competing interests*: The authors declare no competing interests. Correspondence and requests for materials should be addressed to Luc H. Arnal.

