## Supplemental Figure for "Salient 40 Hz sounds probe affective aversion and neural excitability"

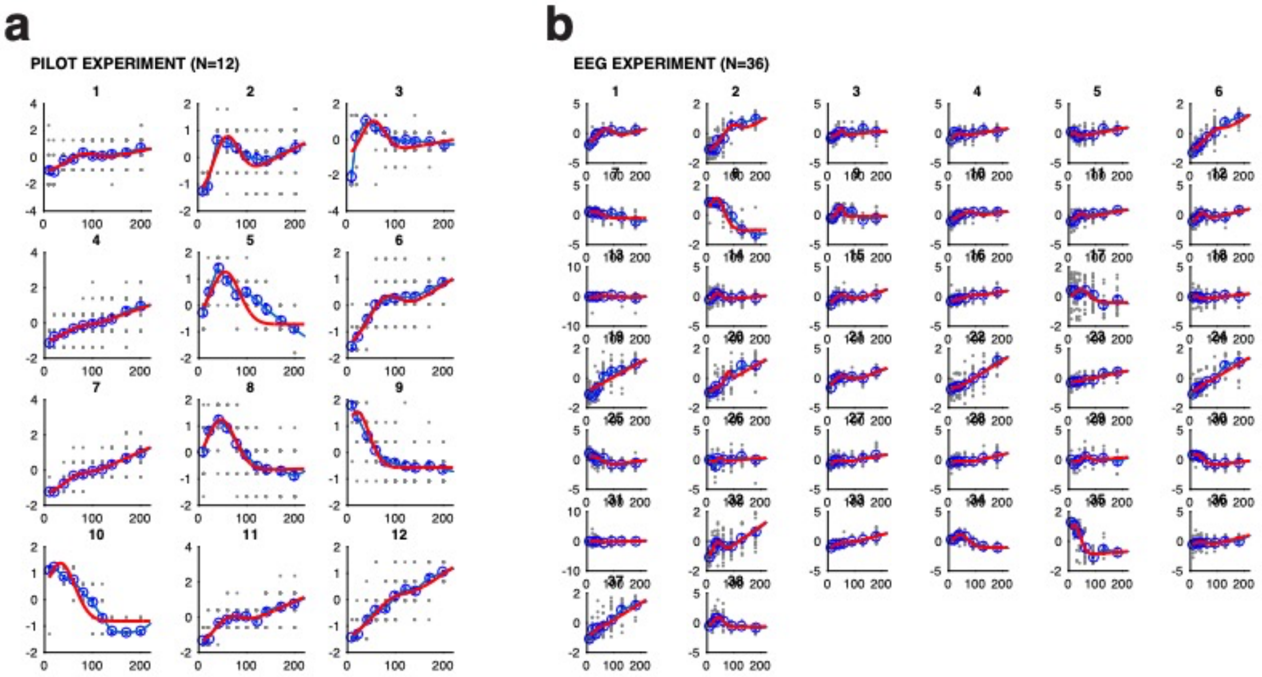


**Non-linear fitting of individual behavioral aversion ratings**. A qualitative inspection of individual aversion profiles suggests aversion is not a linear function of stimulus frequency. While some participants seem more sensitive to low frequencies (*low, 30–60 Hz, e.g.*, participant 5 in Exp. 1) others are more driven by highest frequencies (*high*, >130 Hz, e*.g.*, participant 4 in Exp.1). This suggests that aversion profiles reflect the superposition of two distinct aversion profiles that is determined by individual's relative sensitivity to roughness or pitch: on the one hand, *roughness* responders exhibiting a non-linear response with heightened aversion to low frequencies (*low, 30–60 Hz*), and on the other hand, *pitch* responders, in whom aversion follows a linear profile with a maximal response for the highest frequencies (*high*, >130 Hz). Stemming from the hypothesis that non-linear profiles result from the differential weighting of non-linear *roughness* and *pitch* profiles we then sought to fit individual behavioral data aversion scores. In each participant, we fit a summation between a gaussian distribution and a linear model (see methods). The parameters were fit by minimizing the squared error between the model predictions and the true sound rating taking the stimulation frequency as input. This model allowed us to interpolate our results to untested frequencies and to pool and compare the data across experiments using different stimulus frequencies.
